# The RNA m6A binding protein YTHDF2 promotes the B cell to plasma cell transition

**DOI:** 10.1101/2021.07.21.453193

**Authors:** David J. Turner, Alexander Saveliev, Fiamma Salerno, Louise S. Matheson, Michael Screen, Hannah Lawson, David Wotherspoon, Kamil R. Kranc, Martin Turner

## Abstract

To identify roles of RNA binding proteins (RBPs) in the differentiation of B cells to antibody-secreting plasma cells we performed a CRISPR/Cas9 knockout screen of 1213 mouse RBPs for their ability to affect proliferation and/or survival, and the emergence of differentiated CD138+ cells *in vitro*. We identified RBPs that promoted the appearance of CD138+ cells including CSDE1 and STRAP, as well as RBPs that inhibited CD138+ cell appearance such as EIF3 subunits EIF3K and EIF3L. Furthermore, we identified RBPs that share the property of recruiting the CCR4-NOT complex to their target transcripts have the potential to mediate opposing outcomes on B cell differentiation. One such RBP, the m^6^A binding protein YTHDF2 promotes the appearance of CD138+ cells *in vitro*. In chimeric mouse models YTHDF2-deficient B cells formed germinal centers in a cell-autonomous manner, however plasma cells failed to accumulate.

## Introduction

Humoral immunity requires the generation of antibody-secreting plasmablasts and plasma cells. These are derived from naïve B cell precursors that enter the extrafollicular response and the germinal centre (GC) reaction^1^. Germinal centre derived antibody secreting cells may migrate to the bone marrow where they survive as long-lived plasma cells secreting high-affinity antibody.

The B- to plasma-cell transition requires substantial reprogramming of the transcriptome. This is achieved in part by the actions of transcription factors and epigenetic regulators that act within a layered system to control the timing of differentiation and its coordination with extracellular cues. Dynamic control within this system is further endowed by the integration of signal transduction pathways and post-transcriptional regulators. A central role for post-transcriptional control is mediated by microRNAs such as miRNA148a^2^ and miRNA155^3^ which promote and miRNA125b^4^ which suppresses plasma cell differentiation. RNA binding proteins (RBPs) act co- or post-transcriptionally to influence the quality and quantity of expressed genes. However, while estimates of the numbers of RBP encoded in mammalian genome vary between one-five thousand, few have been implicated in plasma cell differentiation. These include, HNRNPLL which promotes differentiation indirectly by suppressing BCL6 expression^5^; the non-canonical poly(A) polymerase FAM46C/TENT5C which acts as a tumour suppressor in myeloma^6^; and ZFP36L1 which is required for the emigration and survival of plasma cells^7^.

Motivated by a lack of known roles of RBPs in B cell differentiation we developed a CRISPR library targeting mouse RBPs and performed genetic screens for roles in B cell proliferation and survival, and for their differentiation into plasma cells. We identified 103 RBPs that promote and 189 RBPs that inhibit differentiation, and demonstrated YTHDF2, an N6-methyladenosine (m^6^A) binding protein, to be necessary for the accumulation of plasma cells in the bone marrow. Furthermore, we found that RBPs recruiting transcripts to the CCR4-NOT complex for deadenylation and subsequent decay can both promote and inhibit differentiation. Thus, the CCR4-NOT complex is utilised by different adapters to both limit and to promote terminal differentiation.

## Results

### Genetic screening identifies RBPs regulating plasma cell differentiation

We curated a list of 1213 mouse RBPs (excluding ribosomal subunit proteins) compiled from the compendium by Gerstberger et al and presence in at least two RNA interactome capture studies^8–14^ (Supplementary Table 1). A custom sgRNA library targeting these was constructed in a vector backbone expressing puromycin resistance and CD90.1 as selection markers (Supplementary Figure 1A). The library encodes 10 sgRNAs per gene, targets 72 positive control genes and contains 500 negative control sgRNAs. It contains 13,350 sgRNAs in total with >98.8% of sgRNAs normally distributed within a 16-fold range (Supplementary Figure 1B; Supplementary Table 2).

To screen RBPs that regulate cell expansion (proliferation or survival), primary mouse B cells expressing Cas9 from the *Rosa26* locus were cultured *in vitro* on fibroblasts expressing CD40ligand and BAFF for four days with IL-4 followed by four days with IL-21^15^. The representation of sgRNAs at day4 and day8 was determined by next generation sequencing (NGS) and compared (Supplementary Figure 1C; Supplementary Table 3). Guides targeting *Trp53* and *Myc* enriched as expected for inhibiting and supporting cell expansion respectively (Supplementary Figure 1D, E). *Rc3h1, Caprin1* and *Fam46c* were the only RBPs enriched for limiting B cell proliferation or survival (Supplementary Figure 1D, E). *Fam46C* has been implicated in impairing the proliferation and/or survival of B cells^6,16^. Further supporting the predictive power of our dataset, we validated the role for *Rc3h1* as limiting proliferation or survival with individual sgRNAs (Supplementary Figure 1F).

On day8 of the culture we sorted cells based on expression of CD138, a marker of plasma cells and determined the representation of sgRNAs by NGS (Supplementary Figure 1C; Supplementary Table 4). To identify RBPs that regulate differentiation, but not expansion between day 4 and 8, we considered the intersection between the two screens (Figure 1A) and focused on RBPs that enriched for a role in differentiation without enrichment for a role in expansion. The transcription factors: *Bach2, Bcl6, Spi1*, and *Irf8* enriched for inhibiting differentiation, and *Prdm1 and Irf4* enriched for promoting differentiation (Figure 1A, B). These results are consistent with published roles^10–20^ in broadly defining the gene expression profiles of B cells and plasma cells, and therefore underpin our confidence that novel findings in this system reflect meaningful biological effects.

**Fig. 1.**
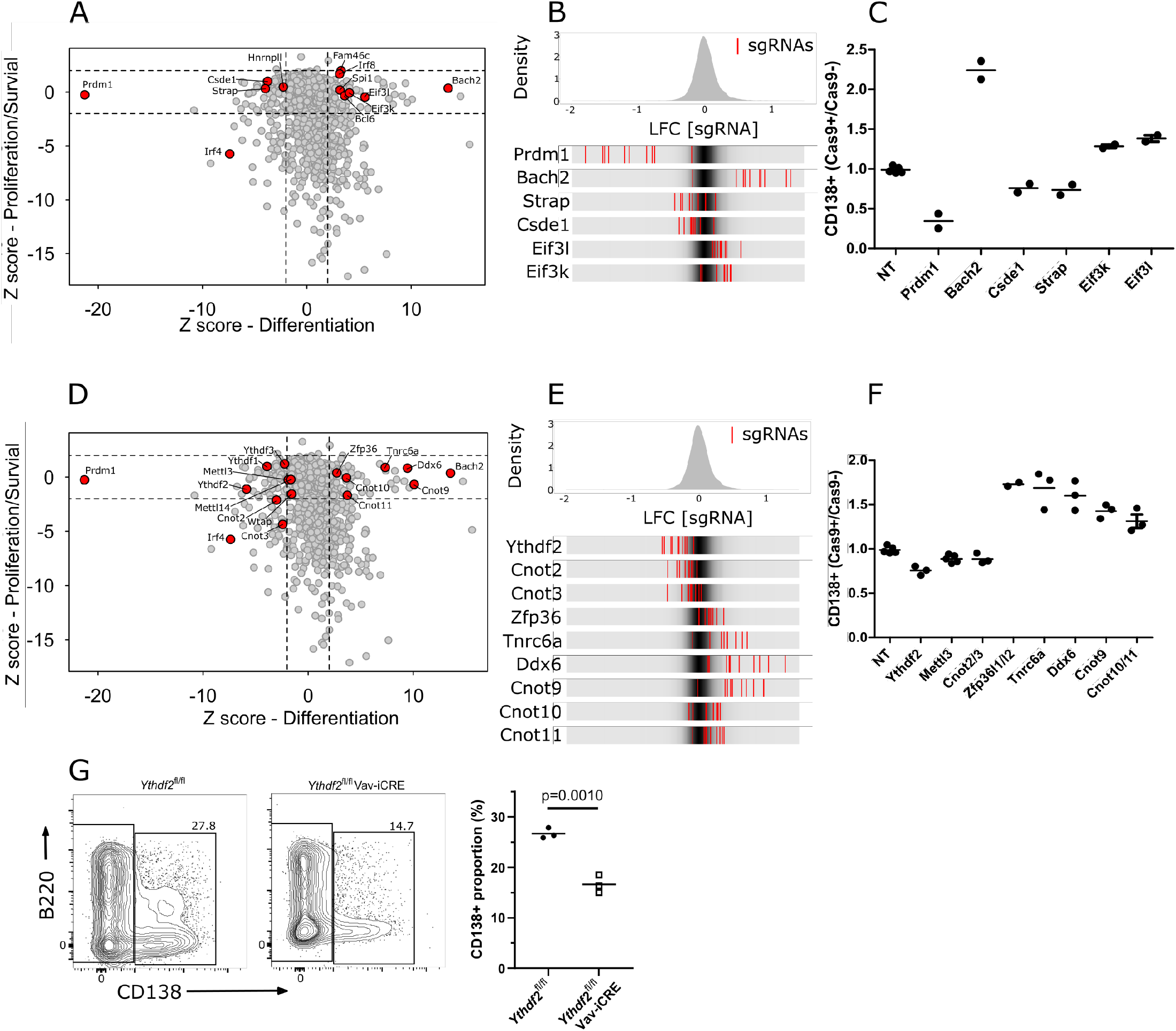
Genetic screen identifies modulators of the B cell to plasma cell transition. **(A)** Dot plot representation of genetic screens of B cell proliferation/survival, and B cell differentiation: X-axis shows z-score of gene-level log_2_ fold change (LFC) for differentiation (CD138+ v CD138- cells) and Y-axis shows z-score of gene-level LFC (day8 v day4 cells) for proliferation/survival calculated by MAGECK. **(B)** Top: Distribution of z-scores of sgRNA-level LFC (CD138+ v CD138- cells) for the differentiation screen. Bottom: LFC (CD138+ v CD138- cells) for all ten sgRNAs targeting the indicated genes. The grey density in the middle of each bar depicts the overall distribution of sgRNAs. Red lines represent individual sgRNAs targeting a given gene. **(C)** The ratio of proportion of cells expressing CD138 in Cas9+ cells and Cas9- cells transduced by viruses with non-targeting (NT), *Prmd1, Bach2, Csde1, Strap, Eif3k, Eif3l* targeting guides at day-8 of the *in vitro* B cell culture. Each symbol is representative of a distinct sgRNA. **(D)** Same as in (A) with additional genes highlighted. **(E)** Same as in (B) with additional genes highlighted. **(F)** Same as in (C) with *Ythdf2, Tnrc6a, Ddx6, Cnot9* targeting guides, with an individual sgRNA that targets *Zfp36l1* and *Zfp36l2*, and paired sgRNAs against *Cnot2*/*Cnot3* or *Cnot10*/*Cnot11*. **(G)** Left: representative flow cytometry for *Ythdf2*^CTL^ and *Ythdf2*^CKO^ and Right: summary data of the proportion of cells expressing CD138 at day 8 of an *in vitro* culture of B cells from *Ythdf2*^fl/fl^ mice (closed circles) or *Ythdf2*^fl/fl^-Vav-iCre mice (open squares). Statistical significance was determined by two-tailed unpaired Student’s t-test. Each symbol is representative of cells from a single mouse.

137 RBPs enriched for a role in plasma cell differentiation that did not enrich for a role in proliferation and/or survival (Supplementary Table 5). We validated CSDE1 and STRAP which bind to each other^28^ as promoting differentiation and the eIF3 octamer subunits eIF3K and eIF3L as inhibiting differentiation (Figure1C). Loss of function mutations in eIF3K and eIF3L have previously been shown to extend *Caenorhabditis elegans* lifespan by enhancing resistance to endoplasmic reticulum (ER) stress^29^. Hence, eIF3K and eIF3L may inhibit CD138+ cell appearance by limiting tolerance to ER stress. All three YTHDF-family members of m^6^A binding proteins were also enriched for promoting differentiation (Figure 1D) of which *Ythdf2* had the most pronounced effect (Figure 1E, F; Supplementary Figure 1G, H). In further support of a role of m6A in promoting B cell terminal differentiation, the three core components of the m6A writer complex (*Mettl3, Mettl14* and *Wtap*), had convergent enrichment on the edge of statistical significance for a role in promoting differentiation (Figure 1D and 1F). By contrast, the ZFP36 family of AU-rich element binding proteins and TNRC6 family critical for miRNA mediated repression were enriched for inhibiting differentiation (Figure1D-F and Supplementary Figure 1G). ZFP36 has been previously reported to limit CD138+ cell appearance *in vitro*^30^.

The YTHDF-, ZFP36- and TNRC6-families of RBPs that are all known to limit target transcript stability by recruiting the CCR4-NOT complex leading to translational repression, deadenylation and subsequent decay^31–33^ show opposing roles in differentiation to CD138+ cells. Furthermore, we found that components of the CCR4-NOT complex had also opposing enrichments: interacting partners CNOT2 and CNOT3 enriched for promoting differentiation, while CNOT9, and the interacting partners CNOT-10 and -11 enriched for limiting differentiation (Figure1D, E). We hypothesise that the CCR4-NOT complex via its interactions with different adapters may act as a post-transcriptional hub that controls the decision to differentiate. In this model, the ZFP36- and TNRC6-families suppress transcripts that promote plasma cell differentiation, while the YTHDF-family enable the transition to plasma cells by suppressing transcripts characteristic of B cells.

### Antibody secreting cells deficient for YTHDF2 fail to accumulate

Our screen demonstrated that *Ythdf2* and its paralogues promote B cell differentiation, without appreciably affecting their proliferation or survival. To confirm this, we took advantage of *Ythdf2*^fl/fl^;*Vav-iCre* (*Ythdf2*^CKO^) mice, in which *Ythdf2* is conditionally deleted from the haematopoietic system^34,35^. Following anti-IgM stimulation *in vitro* the numbers of *Ythdf2*^CKO^ B cells at each division was not different from *Ythdf2*^fl/fl^ control (*Ythdf2*^CTL^) B cells (Supplementary Figure 1I). Moreover, *Ythdf2*^CKO^ B cells showed no difference in the proportions of IgG1+ and IgE+ cells generated *in vitro* compared to *Ythdf2*^CTL^ B cells (Supplementary Figure 1J), indicating that class switch recombination (CSR) did not require YTHDF2 *in vitro*. Because *Ythdf2*^CKO^ B cells had a reduced capacity to differentiate to CD138+ cells compared to *Ythdf2*^CTL^ B cells (Figure 1G), we hypothesise that YTHDF2 promotes B cell differentiation independently of a role in CSR, cell proliferation or survival.

**Supplementary Figure 1.**
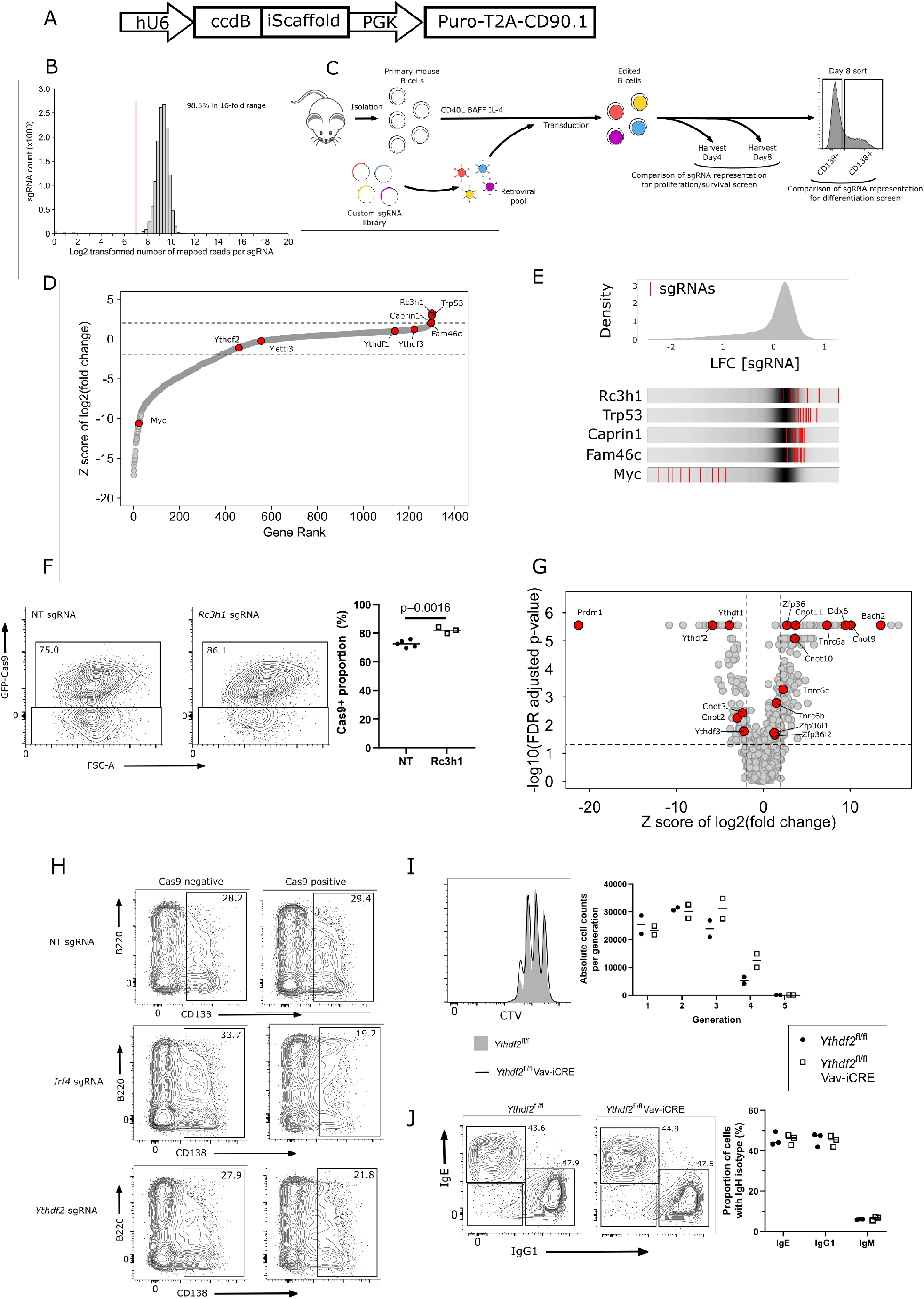
Custom sgRNA library targeting RBPs identifies regulators of B cell proliferation and survival. **(A)** Schematic representation of custom sgRNA library vector backbone. **(B)** Distribution of the relative representation of sgRNAs within the sgRNA library. Red vertical lines indicate the upper and lower bounds of a 16-fold range in representation that includes 98.8% of all sgRNAs. **(C)** Schematic representation of the pooled CRISPR/Cas9 knockout screens in primary mouse B cells. Comparison of sgRNA representation between day4 and day8 identified genes involved in proliferation and/or survival. Comparison of representation between day 8 cells sorted as CD138+ve or -ve identified genes promoting or inhibiting plasma cell differentiation. **(D)** Enrichment plot for the proliferation and/or survival screen. The Y-axis shows z-score of gene-level log_2_ fold change (LFC) (day8 v day4 cells). X-axis shows rank of genes by z-score of gene-level LFC. **(E)** Top: Distribution of z-scores of sgRNA-level LFC for proliferation and/or survival screen. Bottom: LFC (day8 v day4 cells) for all ten sgRNAs targeting the indicated genes. The grey gradient in the middle of each bar depicts the overall distribution. Red lines represent individual sgRNAs targeting a given gene. **(F)** Left: representative flow cytometry and Right: quantification of the proportion of cells in coculture that are expressing Cas9-GFP at day 8 of an *in vitro* culture. Each symbol represents an individual non-targeting (NT) sgRNA (closed circles) or *Rc3h1* sgRNA (open squares). Statistical significance was determined by two-tailed unpaired Student’s t-test. **(G)** Volcano plot representation of CRISPR/Cas9 screen of B cell terminal differentiation X-axis shows z-score of gene-level log_2_ fold change (LFC) (CD138+ v CD138- cells) and the Y-axis shows FDR adjusted p-value calculated by MAGECK. **(H)** Representative flow cytometry showing B220 and CD138 staining of GFP^+^ (Cas9 expressing) and GFP^-^ (Cas9 non-expressing) cells at day-8 of an *in vitro* culture following transduction with viruses expressing non-targeting sgRNA or sgRNA targeting *Ythdf2* and *Irf4*. **(I)** Left: representative flow cytometry analysis of cell trace dilution of 72-hour anti-IgM stimulated B cells. Right: Enumeration of absolute cell number per generation. **(J)** Left: representative flow cytometry analysis for *Ythdf2*^CTL^ and *Ythdf2*^CKO^ of the proportions of cells expressing IgG1 and IgE at day8 of an *in vitro* culture. Right: Quantification of the proportion of cells expressing each IgH isotype.

To investigate an *in vivo* role of *Ythdf2*, we established mixed bone-marrow chimeras in B6.SJL recipients with µMT and either *Ythdf2*^CTL^ or *Ythdf2*^CKO^ bone marrow cells. In these chimeras, CD45.1+ µMT stem cells, which cannot generate B cells, ensure that non-B cells are primarily *Ythdf2*-sufficent, whereas all B cells derive from CD45.2+ control or experimental donors. *Ythdf2*-deficient cells efficiently reconstituted the splenic B cell pool (Figure 2A) enabling us to test the hapten-specific responses of mature B cells lacking YTHDF2 following immunisation with alum precipitated 4-Hydroxy-3-nitrophenylacetyl coupled to the carrier protein Keyhole-Limpet Hemocyanin (NP-KLH). 21 days after immunisation the titer of IgG1 antibodies binding to low-valency hapten was 1.5-fold lower in *Ythdf2*^CKO^ chimeras compared to controls (Figure 2B). In the spleen the number of antigen-specific (NIP+) IgG1+ GC B cells was slightly lower in *Ythdf2*^CKO^ chimeras compared to the controls (Figure 2C). A two-fold reduction in the proportion of GC B cells was previously observed at day14 after NP-KLH immunisation of CD23-CRE *Ythdf2*^fl/fl^ mice, however, their absolute numbers were not reported^36^. In our chimeras, the number of NIP+ IgG1+ splenic plasmablasts (Figure 2D) and NIP+ IgG1+ plasma cells in the bone marrow (Figure 2E) were both three-fold lower in the *Ythdf2*^CKO^ compared to the controls. Together these results demonstrate a requirement for YTHDF2 in the differentiation of antibody secreting cells *in vivo*.

**Figure 2.**
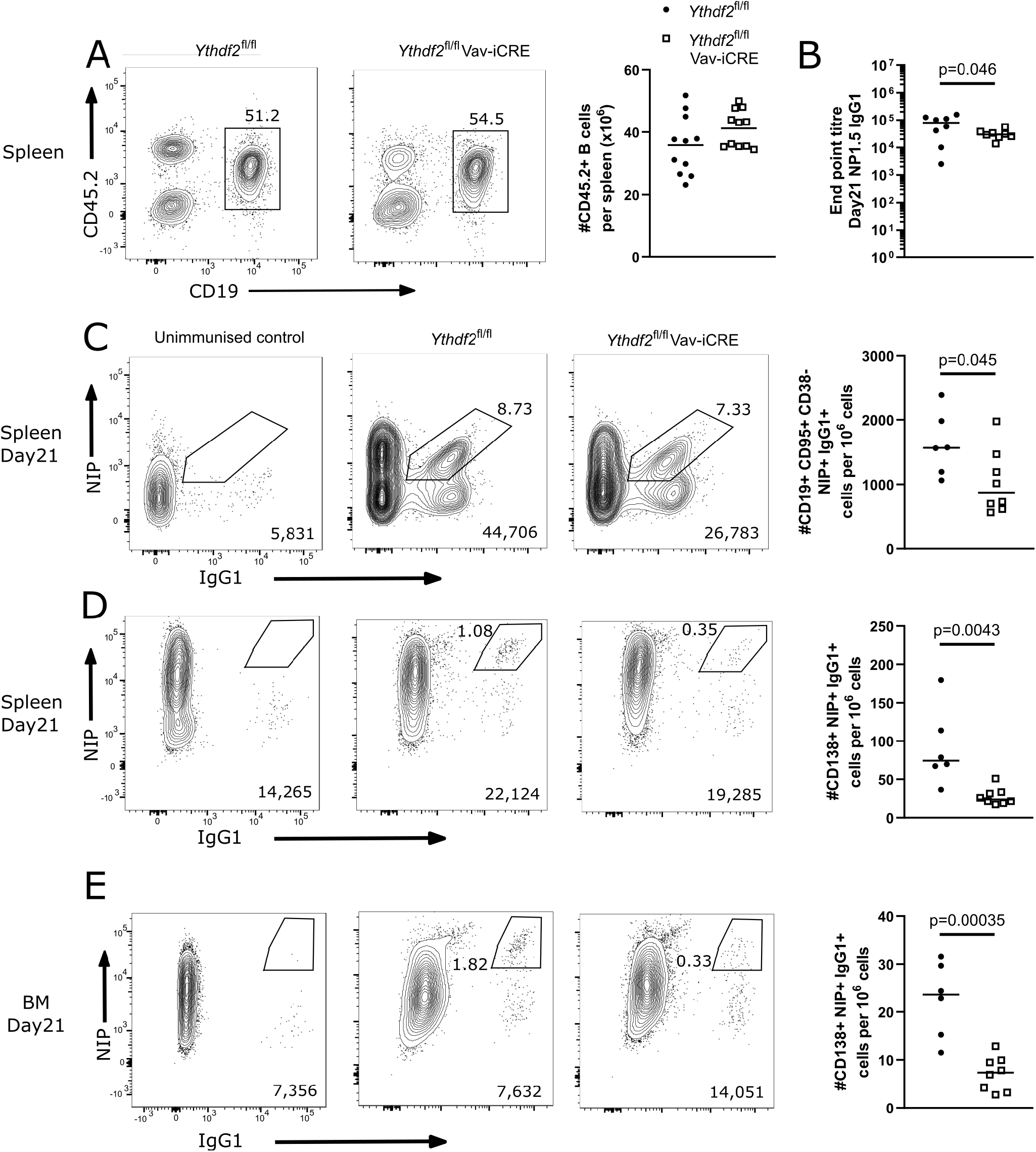
Analysis of µMT chimeras at day 21 after immunization with NP-KLH in alum. **(A)** Left: Representative flow cytometry analysis of cells expressing CD45.2 and CD19 in spleen of µMT chimeras fourteen weeks after reconstitution. Numbers refer to the proportion of viable single cells within CD45.2+ CD19+ gate. Right: The number of cells expressing CD45.2 and CD19. **(B)** End point titre of serum anti-NP1.5 IgG1 antibody. **(C)** Representative flow cytometry analysis of NIP and IgG1 staining on spleen cells gated as CD19+, CD95+ and lacking CD38. **(D)** Representative flow cytometry analysis of NIP and IgG1 intracellular staining of spleen cells gated as CD138+, CD267+ cells; and **(E)** bone marrow CD138+ CD267+ cells. The numbers in the bottom right corner of left-hand panels of B, D and E show the number of events plotted; the numbers adjacent to the gates indicate the proportion of NIP+ IgG1+ cells. The right-hand panels of B, D and E show the number of cells expressing NIP and IgG1 per million viable cells as calculated form the indicated gates. Each symbol represents an individual *Ythdf2*^fl/fl^ control (closed circles) or *Ythdf2*^fl/fl^-Vav-iCre knockout (open squares) mouse. Statistical significance was determined by a two-tailed unpaired Student’s t-test.

To further investigate the role of YTHDF2 in B cell differentiation, we generated competitive mixed bone marrow chimeras in lethally irradiated B6.SJL recipients with equal amounts of CD45.1+ B6.SJL and either CD45.2+ *Ythdf2*^CTL^ or *Ythdf2*^CKO^ bone marrow cells. This system allows YTHDF2-deficient B cells to be in competition with YTHDF2-sufficient B cells at all developmental stages, thus offering a stringent test for functionality of YTHDF2 deficient cells. The *Ythdf2* conditional allele was engineered to express a GFP-YTHDF2 fusion protein in the absence of Cre recombinase, so that GFP expression reliably indicates cells that express YTHDF2^37^. Previously, *Ythdf2* was reported to have a minor role in promoting B cell development^38^. In our competitive chimeras, YTHDF2-deficient cells efficiently reconstituted the mature B cell pool (Figure 3A) with >98% being GFP negative and thus having undergone Cre-mediated deletion (Supplementary Figure 2A). In the spleen, seven days after immunisation with NP-KLH in alum the number of NIP+ IgG1+ GC B cells (Figure 3B) and NIP+ IgG1+ plasmablasts (Figure 3C) were not significantly different between the genotypes. These compartments also showed no evidence of positive selection of non-deleted cells (Supplementary Figure 2A). This indicates that B cells have no essential requirement for *Ythdf2* in the early stages of activation or the extrafollicular response. At 21 days after immunisation the number of antigen-specific class switched GC B cells in the spleen was not appreciably different in *Ythdf2*^CKO^ chimeras compared to the controls (Figure 3D). However, the number of NIP+ IgG1+ splenic plasmablasts was five-fold lower in the *Ythdf2*^CKO^ chimeras compared to the controls (Figure 3E). *Ythdf2*^CKO^ plasma cells were depleted in the bone marrow compartment (Figure 3F), those present were predominantly *Ythdf2* mutants (Supplementary Figure 2B). Moreover, *Ythdf2*^CKO^ antigen specific IgG1+ plasma cells in the bone marrow were also five-fold less abundant in the *Ythdf2*^CKO^ chimeras compared to the controls (Figure 3G). This defect was specific to the mutation of *Ythdf2* as there were similar numbers of CD45.1 B6.SJL derived plasma cells (Supplementary Figure 2C) and NIP+ IgG1+ plasma cells (Supplementary Figure 2D) in the bone marrow of *Ythdf2*^CTL^ and *Ythdf2*^CKO^ chimeras. Taken together, these findings suggest *Ythdf2* is largely dispensable in B cells for activation, the early formation of splenic plasmablasts and participation in the GC response and support a specific role for *Ythdf2* in the differentiation of antibody secreting cells that populate the bone marrow.

**Figure 3.**
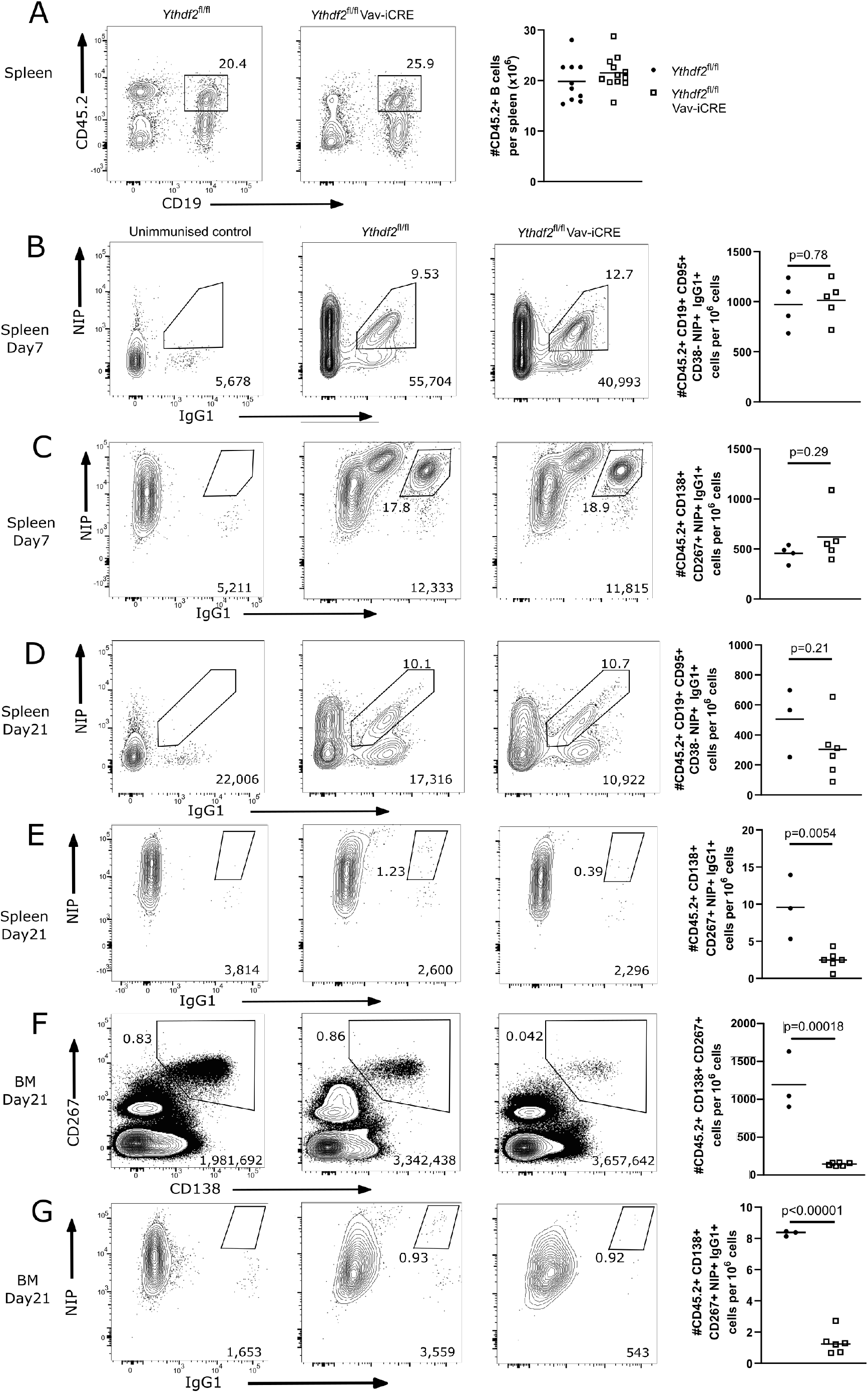
YTHDF2 deficient plasma cells fail to accumulate in the bone marrow. **(A)** Left: Representative flow cytometry analysis of cells expressing CD45.2 and CD19 in spleen of competitive chimeras twelve weeks after reconstitution. Numbers refer to the proportion of viable single cells within CD45.2+ CD19+ gate. Right: The number of cells expressing CD45.2 and CD19. **(B-E)** Representative flow cytometry analysis of NIP and IgG1 staining on spleen cells gated as CD45.2+ CD19+, CD95+ and CD38-ve at day 7 (**B**) and day 21 (**D**), or gated as CD45.2+ CD138+, CD267+ cells at day 7 (**C**) and day 21 (**E**). **(F)** Representative flow cytometry analysis of CD138 and CD267 staining on bone marrow cells gated as CD45.2+ at day 21. (**G**) Representative flow cytometry analysis of NIP and IgG1 staining on bone marrow cells gated as CD45.2+ CD138+, CD267+ at day 21. For each condition, the numbers in the bottom right corner of left-hand panels show the number of events plotted; the numbers adjacent to the gates indicate the proportion of cells within the gate. For each condition, the right-hand plots show the number of cells per million viable cells. Symbols represent data from an individual *Ythdf2*^fl/fl^ control (closed circles) and *Ythdf2*^fl/fl^-Vav-iCre knockout (open squares) mouse. Statistical significance was determined by two-tailed unpaired Student’s t-test.

### The spectrum of m^6^A modified transcripts in B cells

To further understand how the m^6^A modification of mRNA may impact B cell differentiation, we performed m^6^A-eCLIP to identify methylated transcripts within the B cell transcriptome. In total 3,370 high confidence m^6^A sites were identified among 2,658 transcripts in wild-type B cells cultured on 40LB cells for eight days. These m^6^A sites were strongly enriched for DRACH (D=G/A/U, R=G/A, H=A/U/C) and more specifically the RRACU motifs (Figure 4A). m6A sites were enriched in the terminal exon and in the location of the STOP codon in accordance with published datasets (Figure 4B)^39–41^. We hypothesised that m^6^A may mark B cell specific transcripts for turnover upon differentiation. To investigate this, we performed RNA-seq of wild-type B cells cultured on 40LB cells *in vitro* for eight days and sorted by flow cytometry based on B220hi CD138- or B220lo CD138+ expression. However, the m^6^A sites present in the B cell transcriptome (at the 3’UTR and/or CDS) were not characteristic of transcripts specific to either differentiation state and were not biased to transcripts of high expression levels (Figure 4C). As YTHDF paralogues are expected to bind all available m^6^A sites in a transcriptome we speculate that YTHDF2 binds transcripts specific to both differentiation states^42^. However, we are cautious in this because we have not ruled out the possibility that that YTHDF2 has an m^6^A independent role. Among the methylated transcripts identified were *Bach2, Pax5, Irf8* and *Spi1* all of which encode transcription factors that inhibit B cell terminal differentiation, and *Prdm1* which promotes differentiation (Figure 4D). Mechanistically, YTHDF2 promotes the B cell to plasma cell transition, and this function is likely actioned via many transcripts that inhibit this differentiation process.

**Figure 4.**
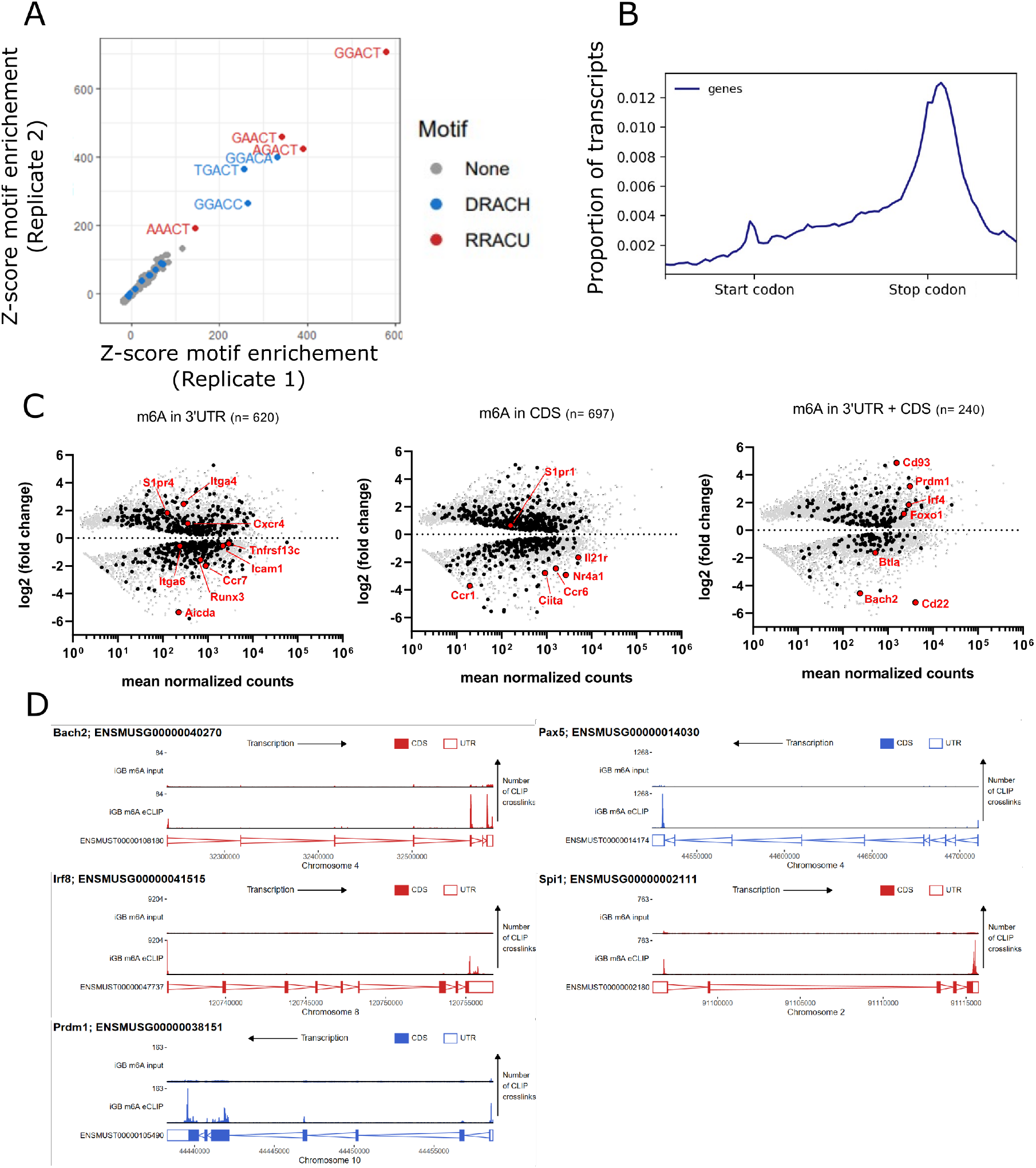
Negative regulators of B cell differentiation are methylated. **(A)** Z-scores showing enrichment of five base motifs centred at *in vitro* derived B cell m^6^A-eCLIP crosslink sites relative to randomised control sites in each replicate. **(B)** Metagene analysis of the location of m^6^A clusters throughout the B cell transcriptome. **(C)** Differential expression (FDR-adjusted p values < 0.05) of methylated (black - identified using the CLIPper pipeline) and unmethylated (grey) transcripts between CD138-B220high and CD138+ B220low *in vitro* cultured cells. (Left) 3’
sUTR methylated transcripts, (centre) CDS methylated transcripts, (right) transcripts methylated in both their 3’UTR and CDS. **(D)** Methylation pattern of individual transcripts encoding regulators (*Bach2, Pax5, Irf8, Spi1, Prdm1*) of B cell differentiation.

In summary, we have evidence that hundreds of RBPs regulate the B cell to plasma cell transition. Among these were both adapters and components of the CCR4-NOT complex. The YTHDF paralogues and the interacting partners CNOT2 and CNOT3 have roles in promoting the appearance of CD138+ cells. The essential role of YTHDF2 is to promote the accumulation of plasma cells in the spleen and bone marrow at the late stages of a response. By contrast, the ZFP36- and TNRC6-families, as well as the CNOT9 and interacting partners CNOT10 and CNOT11 of the CCR4-NOT complex have roles in limiting the appearance of CD138+ cells. The mechanisms by which these RBP limit CD138+ cells remain to be investigated.

## Materials and Methods

### Construction of sgRNA vector backbone and mouse RBP sgRNA library

MSCV_hU6_BbsI-ccdB-BbsI_iScaffold_mPGK_puro-2A-CD90.1 was generated in one round of cloning from a similar plasmid previously generated in our lab^43^ (iScaffold refers to an improved sgRNA scaffold design^44^). A PCR product encoding hU6_BbsI-ccdB-BbsI_iScaffold_mPGK_puro-2A-CD90.1 and the appropriate flanking sequences was generated and ligated by Gibson assembly into a XhoI + SalI linearised vector (MIGRI). Our mouse RBP sgRNA library was generated as previously described^43^. For individual sgRNAs, two 24nt oligonucleotides were annealed and ligated by T4 ligation into our BbsI linearised MSCV backbone vector. For pairwise sgRNAs, PCR products encoding hU6_sgRNA1 and mU6_sgRNA2 with the appropriate flanking sequences were generated and ligated by Gibson assembly into a BbsI linearised RetroQ_BbsI-ccdB-BbsI_iScaffold_mPGK_CD90.1 vector. We targeted mouse RBPs identified by multiple high-throughput studies employing mRNA interactome capture (RIC)^8–14^. Our library targets RBPs identified in at least two mouse RIC studies; the mouse orthologues of RBPs identified in at least two human RIC studies; and manually curated RBPs identified in only one RIC study that had “RNA binding” gene ontology or were paralogues of a human RBP orthologue.

**Supplementary Figure 2.**
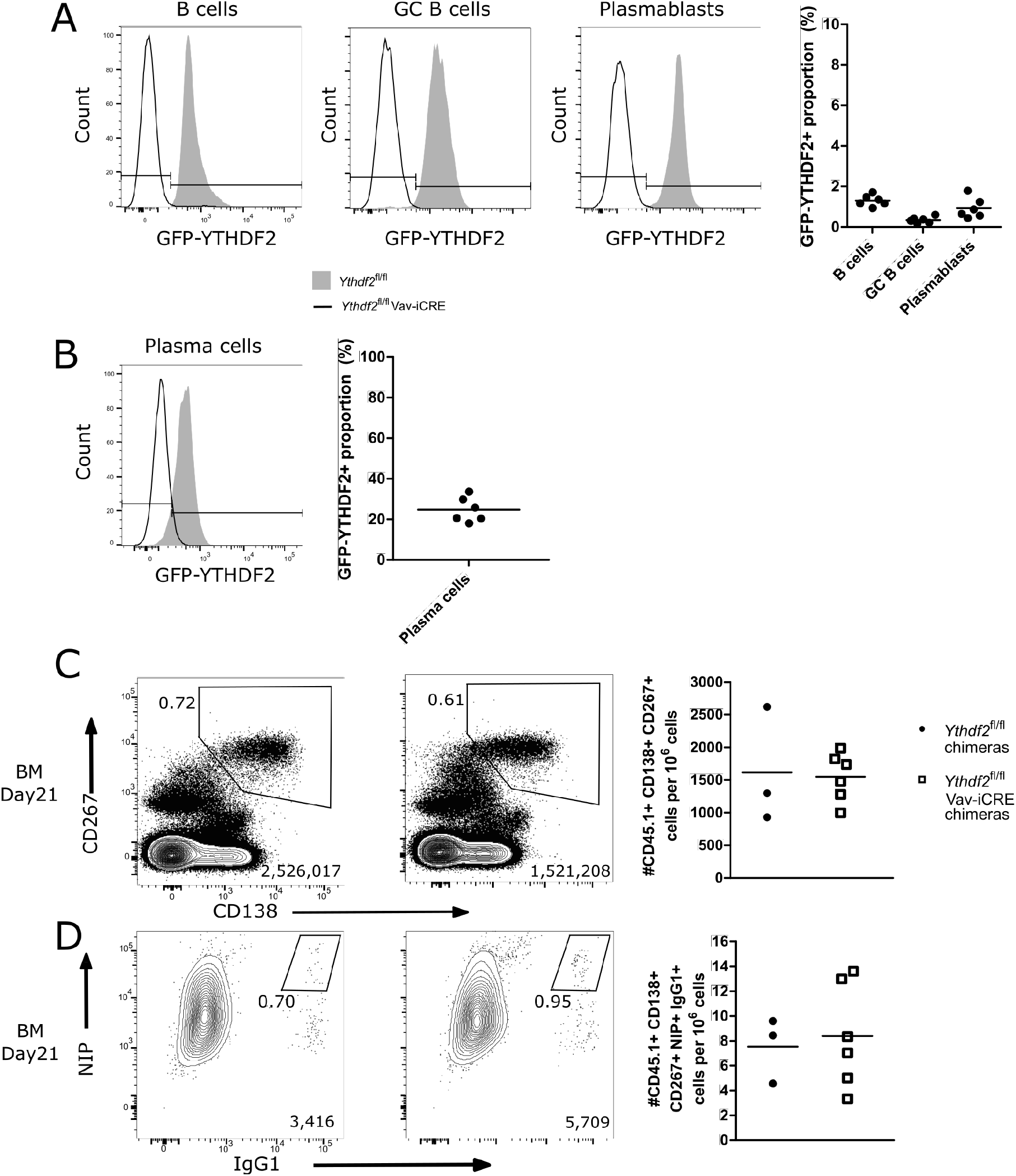
Absence of escapees of *Ythdf2* Cre mediated deletion. **A**) Representative histograms of GFP-YTHDF2 fusion protein expression in surface stained CD45.2 positive total splenic B cells, splenic germinal centre B cells, splenic plasmablasts of competitive chimeras. Right: proportion of CD45.2+ cells within *Ythdf2*^CKO^ competitive chimeras expressing GFP-YTHDF2+. (**B**) Representative histograms of GFP-fluorescence in CD45.2 positive bone marrow plasma cells of competitive chimeras following fixation and intracellular staining for NIP. Right: proportion of CD45.2+ cells within *Ythdf2*^CKO^ competitive chimeras expressing GFP-YTHDF2+. (**C**) Representative flow cytometry analysis of CD138, CD267 staining on bone marrow cells gated as CD45.1+ at day 21. (**D**) Representative flow cytometry analysis of NIP and IgG1 staining on bone marrow cells gated as CD45.21+ CD138+, CD267+ at day 21. For each condition, the numbers in the bottom right corner of left-hand panels show the number of events plotted; the numbers adjacent to the gates indicate the proportion of cells within the gate. For each condition, the right-hand plots show the number of cells per million viable cells. Symbols represent data from an individual *Ythdf2*^fl/fl^ control (closed circles) and *Ythdf2*^fl/fl^-Vav-iCre knockout (open squares) mouse.

### In vitro plasma cell differentiation

3T3 cells expressing mouse CD40L and human BAFF (40LB cells^15^) were maintained in Roswell Park Memorial Institute 1640 (RPMI-1640) media supplemented with 10% FBS and 1x GlutaMAX™ (Gibco: 35050061). Irradiated (∼120Gγ) 40LB cells were seeded at 80×10^3^ cells per well in a 12-well plate in supplemented RPMI-1640 media. The next day, naïve B cells were isolated from the spleens of mice using the B cell isolation kit (Miltenyi Biotec: 130-090-862) and 30×10^3^ cultured on the irradiated 40LB cells with IL-4 in supplemented RPMI-1640 media (final: 10% FBS, 1x GlutaMAX™, 50µM 2-mercaptoethanol Thermo Scientific™: 31350-010, 100 units/ml penicillin, and 100µg/ml streptomycin Thermo Scientific™: 15140, 10ng/ml mIL-4 PeproTech 214-14). On day4 B cells were reseeded on freshly irradiated 40LB cells with IL-21 in supplemented RPMI-1640 media (final: 10% FBS, 1x GlutaMAX™, 50µM 2-mercaptoethanol, 100 units/ml penicillin, and 100µg/ml streptomycin, 10ng/ml mIL-21 PeproTech 210-21). On day8 B cells were harvested and analysed.

### Retrovirus production and transduction

The Plat-E retrovirus packaging cell line was maintained in supplemented Dulbecco’s modified Eagle’s (DMEM) high glucose media (Gibco™: 41965) with 10% fetal bovine serum (FBS). Plat-E cells in logarithmic growth phase were seeded in dishes (Nunc™ 150350). The next day, a transfection mix of 1000µl OptiMEM (Gibco™: 31985), 30µl TransIT®-293 (Mirus® Bio: MIR2700), 9µg transfer vector, 2µg packaging pCL-ECO vector was added dropwise and incubated with the cells overnight. The media was replaced. Then, for two sequential days the viral supernatant was harvested. B cells cultured on 40LBs for three days were transduced with retrovirus particles encoding sgRNAs in the presence of 4ng/ml polybrene (Sigma-Aldrich®: H9268), by spin-fection at 1000g for 45 minutes at 32°C.

### CRISPR/Cas9 genetic screen of B cell differentiation, proliferation and survival

B cells were cultured with 40LB, on day3 cells were transduced with the mouse RBP sgRNA library at an MOI of 0.1 with a predetermined amount of retrovirus supernatant. On day4 transduced cells were positively selected by magnetic assisted cell sorting (MACS) via the cell surface antigen CD90.1 (Thy1.1) using microbeads (Miltenyi Biotec: 130-094-523); the manufacturer’s protocol was adjusted by using 4x less reagents and by maintaining cells at room temperature. On day8, CD138- and CD138+ B cells were physically separated by fluorescence activated cell sorting, washed in PBS, and snap frozen on dry ice in a 15ml conical tube before storage at -80°C for future analysis of terminal differentiation. In addition, on day4 and day8, bulk B cells, unselected for transduction or CD138 expression, were processed and stored in a comparable fashion for future analysis of proliferation/survival. Enough cells to ensure a representation of >1000x transduced cells per sgRNA in the library were maintained at all timepoints and in all populations.

### Next generation sequencing library generation for CRISPR screen

Genomic DNA (gDNA) was isolated as previously described^45^. Next generation sequencing (NGS) libraries were generated as previously described^46^. Multiplexed NGS libraries were sequenced with an Illumina™ HiSeq with a 100bp single end read. The start of the iCRISPR scaffold sequence (GTTTAAGAGCTAT) within each read was identified, and the reads trimmed to encompass the 19 bases immediately preceding this sequence. Bowtie^47^ was used to map these sequences with zero mismatches to a custom genome comprising the sgRNA sequences, and Seqmonk used to quantify the abundance of each sgRNA (https://www.bioinformatics.babraham.ac.uk/projects/seqmonk/). Analysis of our genetic screens was performed with the MAGeCK software^48^. MAGeCK determined gene-level LFC for the proliferation/survival (day8 cells v day4 cells) and differentiation (CD138+ cells v CD138- cells) screens.

### Validation of targets from CRISPR/Cas9 knockout screens

A mixed population of GFP-Cas9 positive and GFP-Cas9 negative B cells, at a ratio of 75:25, were cocultured and transduced with individual sgRNA vectors against genes of interest. The ratio of transduced cells that were differentiated between the Cas9 positive and Cas9 negative populations was calculated and used to control for the variation of the *in vitro* germinal centre B cell culture between experiments. Furthermore, the change in GFP+: GFP-ratio over the culture indicated whether genetically modified cells (Cas9+) had a competitive advantage over unperturbed cells (Cas9-).

### Proliferation assay

Naïve B cell were labelled with CellTrace™ Violet (Thermo Scientific™: C34557), following the manufacturer’s protocol, then seeded at 2×10^6^ cells/ml in 100µl per well of a 96 well plate and stimulated with anti-IgM (2.5µg/ml) (F(ab’)_2_ fragment, polyclonal) (Jackson ImmunoResearch: 115-006-020) with IL-4 (10ng/ml) and IL-21 (10ng/ml) (PeproTech: 210-21. The inclusion of counting beads enabled the calculation of absolute cell numbers.

### Mice

All mice were on a C57BL/6 background. R26-GFP-Cas9 Mb1-Cre mice were derived from C57BL/6^Cd79atm1(cre)Reth 49^ and C57BL/6^Gt(ROSA)26Sortm1(CAG-cas9*,-EGFP)Fezh 50^. For µMT bone marrow chimera experiments B6.SJL-*Ptprc*^*a*^*Pepc*^*b*^/^Boy^ (B6.SJL) mice were used as recipients and reconstituted with CD45.1 µMT (Ighm^tm1Cgn^)^51^ and either *CD45*.*2 Ythdf2*^CTL^ (*Ythdf2*^*Tg*(Vav1-icre)A2Kio^) or *CD45*.*2 Ythdf2*^CKO^ (*Ythdf2*^*tm1*.1Doca + Tg(Vav1-icre)A2Kio^) mice^34,35,37^. For competitive bone marrow chimeras B6.SJL-*Ptprc*^*a*^*Pepc*^*b*^/^Boy^ (B6.SJL) mice were used as recipients and reconstituted with B6.SJL mice and either *CD45*.*2 Ythdf2*^CTL^ or *CD45*.*2 Ythdf2*^CKO^ mice.

All mouse experimentation was approved by the Babraham Institute Animal Welfare and Ethical Review Body. Animal husbandry and experimentation complied with existing European Union and United Kingdom Home Office legislation and local standards. Mice were bred and maintained in the Babraham Institute Biological Support Unit. Since the opening of this barrier facility in 2009 no primary pathogens or additional agents listed in the FELASA recommendations have been confirmed during health monitoring surveys of the stock holding rooms. Ambient temperature was ∼19-21C and relative humidity 52%. Lighting was provided on a 12-hour light: 12-hour dark cycle including 15 min ‘dawn’ and ‘dusk’ periods of subdued lighting. After weaning, mice were transferred to individually ventilated cages with 1-5 mice per cage. Mice were fed CRM (P) VP diet (Special Diet Services) ad libitum and received seeds (e.g. sunflower, millet) at the time of cage-cleaning as part of their environmental enrichment.

### Generation and primary immunisation of chimeric mouse models

B6.SJL recipient mice were lethally irradiated with two doses of 5.0Gy. For µMT chimeras, recipients were reconstituted with 3×10^6^ total bone marrow cells composed from 80% µMT donor bone marrow cells and 20% either *Ythdf2*^CTL^ or *Ythdf2*^CKO^ donor bone marrow cells. For competitive chimeras, recipients were reconstituted with 3×10^6^ total bone marrow cells composed from 50% B6SJL donor bone marrow cells and 50% either *Ythdf2*^CTL^ or *Ythdf2*^CKO^ donor bone marrow cells. Mice received intraperitoneal injection of 200µl sterile PBS containing 100µg of NP(23)KLH adsorbed in 40% v/v Alum (Serva™: 12261). NP-KLH was emulsified in Alum by rotation for 30 minutes at room temperature protected from light.

### ELISA

NP specific antibodies were detected by ELISA as previously described^7^; antibody end point titres were used as a measure of relative concentration.

### Flow cytometry

Cell suspensions from spleen and bone marrow were prepared and stained as previously described^7^. A list of antibodies is provided in Supplementary Table 6.

### m^6^A-eCLIP

B cells were cultured *in vitro* and on day8 total B cell and plasma cells were separated from 40LB cells by negative selection with H-2Kd-biotin antibody and anti-biotin microbeads (Miltenyi 130-090-485) following the manufacturer’s instructions. Total RNA was prepared with a kit (Zymo R2050) following the manufacturer’s instructions and sent to Eclipse Bioinnovations who isolated poly(A) RNA, crosslinked an m^6^A antibody, performed immunoprecipitation and NGS.

Data were analysed by Eclipse Bioinnovations according to their standard m^6^A-eCLIP analysis pipeline (https://eclipsebio.com/wp-content/uploads/2021/06/eclipsebio_data_analysis_review_m^6^A-eCLIP.pdf). Briefly: The first 10 bases of each read (which comprise a UMI) were trimmed using UMI-tools^52^, followed by trimming of poor quality and adaptor sequences from the 3’ end of the reads using cutadapt. Reads mapping to repetitive elements were filtered out, and the remaining reads mapped to the mouse GRCm38 genome build using STAR. PCR duplicates were removed using UMI-tools. Clusters of reads (peaks of m^6^A modification) were identified using CLIPper (https://github.com/YeoLab/clipper/wiki/CLIPper-Home), and log2 fold change relative to the corresponding input sample calculated. Clusters that were reproducible between replicates were then identified using IDR^53^ (https://github.com/nboley/idr). Single nucleotide resolution crosslink sites with enrichment relative to input were also identified using PureCLIP^54^. The gene or feature type to which each crosslink site was assigned was based on the Ensembl Mouse GRCm38.97 annotation release. If the site overlapped multiple genes or feature types, the most likely was chosen in a hierarchical way: first, protein coding isoforms were prioritised; second, transcript isoforms with support level < 4 were prioritised; third by a hierarchy of feature types (CDS > 3’UTR > 5’UTR > intron > non-coding exon > non-coding intron); third, higher confidence isoforms (based first on transcript support level and second on whether they have a CCDS) were prioritised.

To assess enrichment of 5 base motifs at the crosslink sites, the 5 bases centred on each crosslink site for each replicate were identified, and the representation of all 5-mers quantified. This was repeated for 100 sets of control sites, where the locations of the crosslink sites were randomised across all genes that contain at least one crosslink site, ensuring the same distribution across features as in the m^6^A-eCLIP dataset. The z-score for each 5mer was then calculated as: (occurrence at eCLIP crosslink sites - mean occurrence at control sites) / standard deviation of occurrence at control sites.

Metagene analysis of the distribution of m^6^A was performed on the Galaxy server using deepTools^55^. The computeCoverage tool was first used using the Ensembl Mouse GRCm38.97 GTF file, scaling all CDS annotations and additionally analysing 1kb up- and downstream of each. A BED file containing the reproducible clusters was used as the score file, and the maximum value calculated, without skipping 0s (therefore, this will be 1 for windows containing a cluster and 0 for those without). The resulting matrix was then used as input to plotProfile, and the average across all genes plotted.

### Generation and analysis of mRNA sequencing libraries

For transcriptomic analysis, total RNA was isolated from 0.4×10^6^ FACS-sorted B220hi CD138- and B220low CD138+ B cells (derived from 3 biological replicates after eight days in culture) using the RNeasy Mini Kit (Qiagen). cDNA was generated from polyadenylated transcripts employing the SMART-Seq v4 ultra low input RNA kit (Takara Bio). RNA and cDNA quality was analysed on a 2100 Bioanalyser (Agilent). mRNAseq libraries were prepared using Nextera XT DNA library preparation kit (Illumina) and quantified with KAPA library quantification kit (Roche). Barcoded libraries were multiplexed and sequenced on an Illumina HiSeq 2500-RapidRun system on a 50bp single-end mode with a coverage of 20M reads per sample. Reads were trimmed using Trim Galore and mapped to mouse genome GRCm38.97 using HiSat2 (2.1.0)^56^. Raw counts were calculated over mRNA features (excluding Ig molecules) using SeqMonk (1.47.2; https://www.bioinformatics.babraham.ac.uk/projects/seqmonk/). DESeq2 (1.30.1)^57^ was used to calculate differential RNA abundance and performed using default parameters, with “normal” log2 fold change shrinkage. Information on biological replicates were included in the design formula to have paired analysis. Differentially expressed transcripts were cross referenced with transcripts that contained peaks of m^6^A modification identified using CLIPper.

## End Matter

### Author Contributions and Notes

D.T. conceptualization; methodology; investigation; formal analysis; visualization; data curation; writing - original draft preparation - review & editing; project administration. A.S. conceptualization; investigation; methodology; writing - review & editing. F.S. conceptualization; formal analysis; visualisation; writing - review & editing. L.S.M. conceptualization; formal analysis; visualization; data curation; writing - review & editing. M.S. conceptualisation, methodology; validation; writing - review & editing. H.L. and D.W. resources. K.R.K. conceptualization; supervision; funding acquisition; writing - review & editing. M.T. conceptualization; writing - review & editing; project administration; funding acquisition; supervision.

### Materials and data availability

The sgRNA library is available upon request and from Addgene (#169082). The CRISPR/Cas9 knockout screen data and m6A-eCLIP data that support the findings of this study have been deposited in GEO with the GSE179919 accession code, and the RNA-seq data has been deposited in GEO with the GSE179281 accession code.

## Acknowledgments

We thank the Babraham Institute Biological Support Unit, Flow Cytometry and Bioinformatics Facilities for outstanding support, and D. Hodson and C. Ribeiro de Almeida for helpful discussions and comments on the manuscript. This study was supported by funding from the Biotechnology and Biological Sciences Research Council (BBSRC) (BBS/E/B/000C0427; BBS/E/B/000C0428; the BBSRC Core Capability Grant to the Babraham Institute; a BBSRC-iCASE studentship BB/L016745/1 in partnership with Abzena; and a Wellcome Investigator award (200823/Z/16/Z) to M.T. F.S. was supported by European Molecular Biology Organization (EMBO) Long-Term Fellowship (ALTF 880-2018). K.R. K’s laboratory is funded by a Cancer Research UK program grant (C29967/A26787) and project grants from the Medical Research Council, Blood Cancer UK, Barts Charity, and the Kay Kendall Leukaemia Fund. We thank the Finkelstein Lab (https://github.com/finkelsteinlab) for the BioRxiv template.

